# Iron-sensitive RNA regulation by poly C binding proteins

**DOI:** 10.1101/2024.10.17.618301

**Authors:** Kwame Forbes, Grant A. Goda, Grace A. Eramo, Conner Breen, Douglas F. Porter, Paul A. Khavari, Daniel Dominguez, Maria M. Aleman

**Affiliations:** Bioinformatics & Computational Biology, University of North Carolina at Chapel Hill; Department of Pharmacology, University of North Carolina at Chapel Hill, USA; Department of Chemistry, University of North Carolina at Chapel Hill, USA; Program in Epithelial Biology, Stanford University, Stanford, CA, USA; Department of Biochemistry and Biophysics, University of North Carolina at Chapel Hill, USA; Lineberger Comprehensive Cancer Center, University of North Carolina at Chapel Hill, USA; RNA Discovery Center, University of North Carolina at Chapel Hill, USA; Blood Research Center, University of North Carolina at Chapel Hill, USA; McAllister Heart Institute, University of North Carolina at Chapel Hill, USA

## Abstract

Iron is essential for normal cellular function. Homeostatic responses to low iron availability have long been known to rely on posttranscriptional mechanisms. Poly C binding proteins (PCBPs) are essential RNA binding proteins that regulate alternative splicing, translation and RNA stability. They also serve as critical iron chaperones that manage intracellular iron flux. However, the impact of cellular iron levels on the PCBP-directed transcriptome has not been globally evaluated. We have found broad transcriptome changes in response to low iron availability consistent with numerous operant posttranscriptional mechanisms that sense iron. By comparing gene expression and alternative splicing directed by PCBPs to the iron-sensitive transcriptome, we found genes with iron-sensitive PCBP RNA regulation. Further, we demonstrate that cellular iron levels impact PCBP1 RNA binding. Our data show a large-scale RNA regulatory response to cellular iron levels and open the possibility for other iron-sensitive RNA binding proteins.

## INTRODUCTION

Gene expression and metabolism are intertwined processes that both rely on the availability of metal ions such as iron. Iron supports reactions needed to carry out important cellular functions such as DNA synthesis and oxidative phosphorylation and is essential for the activity of iron-dependent enzymes (*e.g.*, demethylases or phosphatases)^1^. Too much or too little iron is problematic for cell function and can lead to toxicity or stress. Thus, organisms have evolved both systemic and cellular mechanisms to control the availability of iron. RNA regulation is a critical component of intracellular iron homeostasis and involves the RNA binding activity of iron regulatory proteins (IRP) 1 and 2. Through specific interactions with structured iron regulatory elements (IREs) in mRNA untranslated regions, IRPs can promote or repress the translation of key iron homeostasis genes. Mechanisms controlling the ability of IRPs to bind RNA are iron-dependent. Thus, when intracellular iron levels are low, IRP-IRE interactions are enhanced leading to increased translation of mRNAs like transferrin receptor 1 (*TFRC*) or decreased translation, like ferritin heavy chain (*FTH1*)^2^. The net effect of this posttranscriptional regulation is to maintain iron levels and enable normal cellular function.

Recently, two additional RNA-binding proteins (RBPs) have been demonstrated as ‘iron-sensitive.’ The first, tristetraprolin (also known as ZFP36) is upregulated transcriptionally during iron deficiency to specifically reduce the stability of mRNAs for iron-dependent proteins in the electron transport chain, consequently driving a shift from oxidative phosphorylation to glycolysis to conserve iron^3, 4^. The second RBP with iron-sensitive activity is SRSF7—a splicing factor that promotes the exclusion of alternative exons. Iron disrupts the RNA binding of the SRSF7 to target exons, including cell death receptor Fas/CD95, leading to differential regulation of alternative splicing^5^. These recent discoveries suggest a broader posttranscriptional response to iron levels than previously appreciated and that genes targeted for iron-sensitive regulation go beyond iron homeostasis genes.

Beyond post-transcriptional control of genes involved in iron import, export, and storage, iron itself is carefully managed in the cell. Protein iron chaperones are used across life forms to metallate specific ferrous iron-dependent proteins or to deliver iron where it is needed intracellularly^6^. Intriguingly, the first mammalian iron chaperone discovered is best known for its well established and canonical role as an RBP. Via a yeast-based screen with a human liver cDNA library, poly(C) binding protein 1 (PCBP1) was discovered to deliver iron to ferritin, the iron storage complex^7^. Likewise, other PCBP family members (PCBP2-4) share iron chaperone activity^8^. The iron chaperone activity of PCBPs is important for managing iron uptake, iron storage, iron-sulfur cluster formation, heme biosynthesis, and the response to hypoxia^7–15^.

Yet, PCBPs are also important regulators of RNA processing, with roles in alternative splicing, RNA stability, and translation^16–19^. PCBPs are broadly expressed, especially PCBP1 and PCBP2, and at relatively high concentrations^20^. Seminal studies have shown that these PCBPs bind thousands of cellular mRNAs (in crosslinking and immunoprecipitation (eCLIP) experiments) and control the RNA processing fate of many genes^21^, indicative of a broad role in gene regulation. Indeed, PCBP1 and PCBP2 are both essential for life^18^.

The intriguing combination of iron and RNA binding by PCBPs raises interesting questions on how these actions may impact overall PCBP function. For example, does iron binding alter RNA binding? Could PCBPs operate similarly to IRPs? While recent work suggests iron binding and RNA binding by PCBPs are both ‘separable and independent’^22^, two prior *in vitro* studies found that iron reduces PCBP RNA binding^23, 24^. Whether these results translate to cellular RNA binding is not yet clear. Given the broad role of RBPs in modulating gene expression, the potential impact of iron-sensitive RNA regulation by PCBPs warrants further investigation; thus, we have undertaken the following study.

Here, we use transcriptomic and in-cell RNA binding approaches to determine how cellular iron levels impact RNA regulation and identify massive scale expression and RNA splicing changes in response to iron chelation. We investigate the involvement of PCBPs in RNA regulatory responses to iron chelation and a role for PCBP1 in iron-sensitive splicing. We further demonstrate PCBP1-RNA interactions in cells are enhanced by iron chelation, suggesting that iron has a direct impact on the way PCBP1 interacts with RNA. With PCBPs functioning as both iron chaperones and RBPs, this family of proteins is well situated to integrate iron sensing with control of the cellular adaptive response to low iron via changes in gene expression.

## RESULTS

### Iron chelation broadly alters the transcriptome

Our overall hypothesis is that PCBPs regulate RNA in an iron-sensitive manner. To explore this, we first evaluated how the transcriptome changed in response to iron chelation. We performed deep mRNAseq on RNA from K562 cells treated overnight with or without a cell-permeable iron chelator (21H7^25^). K562 cells, an erythroid leukemia cell line, were chosen for these experiments due to their known PCBP expression and RNA regulatory activity^26–29^, as well as the public availability of enhanced crosslinking and immunoprecipitation (eCLIP) data from the ENCODE consortium^30^ that identifies K562 PCBP target RNAs. To confirm that the 21H7 iron chelator functioned as expected, we validated known changes in protein levels for ferritin heavy chain (FTH1)^31^ and iron regulatory protein 2 (IRP2)^32^ by western blot. With iron chelation, FTH1 was decreased, while IRP2 was increased (Fig. 1A-C). At the RNA level, we observed large-scale changes in gene expression; 1899 genes were significantly upregulated (*P* <0.05 and log2 fold change > 1), and 586 genes were significantly downregulated (*P* <0.05 and log2 fold change < -1; Fig. 1D and Supplemental Table 1). We evaluated the expression of genes known to change in response to low iron in K562 cells. We observed increased mRNA levels for transferrin (*TF*) and transferrin receptor 1 (TFR1, gene *TFRC*)^33^ and decreased mRNA levels for ferroportin (*SLC40A1*)^34^ (Fig. 1E). Over 500 genes were more upregulated than *TFRC* and over 100 genes were more downregulated than *SLC40A1* which highlights the extent and robustness of gene expression changes in response to low iron availability.

**Figure 1.**
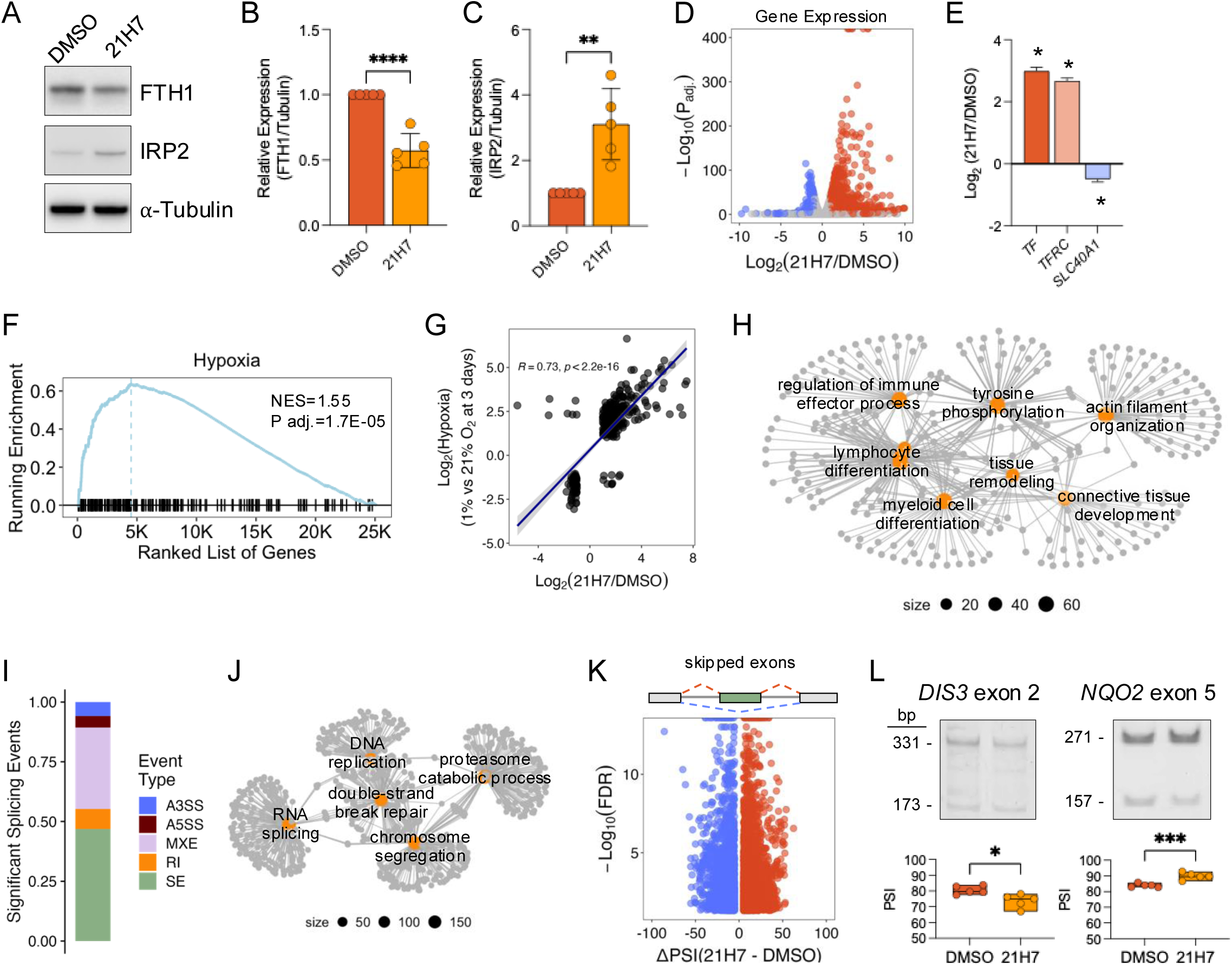
**A.** Western blot analysis FTH1 and IRP2 after treatment of K562 cells with iron chelator, 21H7, or DMSO. Tubulin was used as loading control. **B-C.** Western blot quantification of FTH1 and IRP2, respectively, after treatment. Data are mean ± standard deviation (SD). \**P*<0.05, \*\**P*<0.01. **D.** Volcano plot of differentially expressed genes (DEG) after iron chelation. Blue points represent down-regulated genes, and red points represent up-regulated genes. **E.** Bar plot of key DEGs (*TF, TFRC, SLC40A1*) after iron chelation. Data are mean ± standard error of the mean (SEM). **F.** Enrichment of hallmark hypoxia genes by GSEA after iron chelation. NES, normalized enrichment score. **G.** Scatter plot of DEGs that overlap significantly between iron chelation (21H7 vs. DMSO) and Hypoxia (1% vs. 21% O2 at 3 days){Jain, 2020 #333} with Spearman’s rank correlation. **H.** Network plot of gene ontology (GO) terms from significantly DEGs after iron chelation. GO term nodes are orange and genes in each node are grey circles. **I.** Bar plot of the proportion of significant alternative splicing (AS) events in chelator vs. control for each event type. Skipped exons (SE), alternative 5’ splice site (A5SS), alternative 3’ splice site (A3SS), retained intron (RI), mutually exclusive exon (MXE). **J.** Network plot of GO terms from significant AS genes in chelator vs. control. **K.** Volcano plot of SE after iron chelation. Blue points are excluded exon events, and red points are included exon events. **L.** RT-PCR analysis and quantification for AS events in *DIS3* and *NQO2* mRNA after treatment.

We performed gene set enrichment analysis (GSEA) of hallmark gene sets^35^ and observed significant enrichment of hypoxia genes (Fig. 1F). Iron chelators, such as desferoxamine (DFO), are often used to simulate hypoxia *in vitro* and *in vivo*^36–38^, thus enrichment for this pathway is expected. Indeed, we found a significant positive correlation in differential gene expression between our iron chelation data set and a hypoxia treatment data set^39^ in the same cell line (Fig. 1G). With iron chelation, we also found significant enrichment by GSEA for multiple signaling pathways including TNFα signaling (Supplemental Table 2), consistent with prior work showing TNFα signaling promotes sequestration of iron^40, 41^. Gene ontology analysis of differential genes identified hits amongst lymphoid and myeloid cell differentiation, tyrosine phosphorylation, and actin filament organization (Fig. 1H and Supplemental Table 3), highlighting the broad role of iron (and the regulation of gene expression) in growth and development. Together, these data clearly show pervasive transcriptome-wide changes after modulating the availability of a single ion in cells and changes in expected and previously unappreciated iron-sensitive pathways.

Because changes in differential gene regulation result from multiple layers (*e.g.*, altered transcription or RNA processing), we next evaluated alternative splicing in response to iron chelation. We detected thousands of significant alternative splicing events, including skipped exons, retained introns, mutually exclusive exons, and 5’ and 3’ alternative splice sites (Fig. 1I and Supplemental Fig. S1A-D). Genes affected at the level of splicing were involved in pathways such as DNA replication, chromosome segregation, and RNA splicing (Fig. 1J and Supplemental Table 4). Iron is already well established to directly support DNA replication as a cofactor for ribonucleotide reductase that generates dNTPs^42^, however the role of iron-sensitive splicing in DNA replication has not been shown. Alternative splicing is known to be operant during the cell cycle^43^ and chromosome segregation during mitosis is sensitive to spliceosome dysfunction (reviewed in ^44^). Finally, RNA splicing genes themselves show iron-sensitive regulation possibly feeding forward to the massive scale alterations in splicing we observed (*i.e.*, regulating the regulator). This analysis suggests cell cycle control during iron chelation is greatly affected by changes in alternative splicing.

We next looked further at the most abundant splicing event type, skipped exons, which occur when an exon can be included or excluded (Fig. 1K, top). We measured percent spliced in (PSI) values—the percentage of transcripts for a given gene including a particular exon—after iron chelation and found nearly 5,000 significant skipped exon events (Fig. 1K, bottom). To the best of our knowledge such a dramatic impact of iron chelation on RNA splicing has not been reported. We validated skipped exon events in *DIS3* and *NQO2,* representing both decreased and increased exon inclusion, respectively (Fig. 1L). DIS3 is a component of the RNA exosome with ribonuclease activity involved in cell differentiation pathways^45–47^ and NQO2 is an oxidoreductase involved in metabolism^48, 49^.

Overall, iron chelation had a profound impact on the transcriptome of K562 cells and the abundance of changes both in gene expression and alternative splicing support a model where co-or posttranscriptional RNA regulation is highly sensitive to cellular iron levels.

### PCBP1-directed transcriptome alternations

We reasoned we could detect iron-sensitive PCBP RNA regulatory activity by comparing changes in gene expression and alternative splicing between iron chelation datasets and PCBP knockdown or overexpression datasets. To do this, we performed deep mRNAseq after PCBP1 shRNA-mediated knockdown or inducible PCBP1 overexpression in K562 cells. To knockdown PCBP1, we treated K562 cells with lentiviral particles containing a shRNA gene targeting PCBP1 (shPCBP1) or luciferase-targeting shRNA (shCtrl) (Fig. 2A). Western blotting for PCBP1 confirmed near complete depletion after viral infection and selection (Fig. 2B). We found 296 significantly differentially expressed genes with PCBP1 knockdown, and as expected PCBP1 itself was the most changed, further confirming knockdown (Fig. 2C and Supplemental Table 5). A single hallmark gene set was enriched with PCBP1 knockdown, E2F Targets (Fig. 2D), which includes genes involved in DNA replication and cell cycle control (Supplemental Table 2). To induce PCBP1 overexpression in K562 cells, we first generated K562^flox^ cells using Cas9/sgRNAs targeted to the AAV1 locus and a linearized plasmid containing a blasticidin resistance gene flanked by lox2722 and loxP sites^50^. Subsequently, we used recombinase-mediated cassette exchange (RMCE)^51^ to integrate a cassette containing flag-tagged PCBP1 under the control of a tet-inducible promoter and puromycin resistance (Fig. 2E). After induction with doxycycline, PCBP1 was four-fold overexpressed (Fig. 2F). mRNAseq identified 251 significantly differentially expressed genes with PCBP1 overexpression. As expected, the most significant differentially expressed gene was PCBP1 itself (Fig. 2G and Supplemental Table 6). GSEA found the iron pathway, heme metabolism, significantly enriched after PCBP1 overexpression, while hypoxia was significantly downregulated (Fig. 2H and Supplemental Table 2). These hits with PCBP1 overexpression show a striking connection between PCBP1, iron and gene regulation. Despite the relatively limited set of changes in gene expression for PCBP1 knockdown and overexpression, we observed a significant negative correlation in fold changes between data sets (Fig. 2I), underscoring that these are PCBP1-directed changes in mRNA levels.

**Figure 2.**
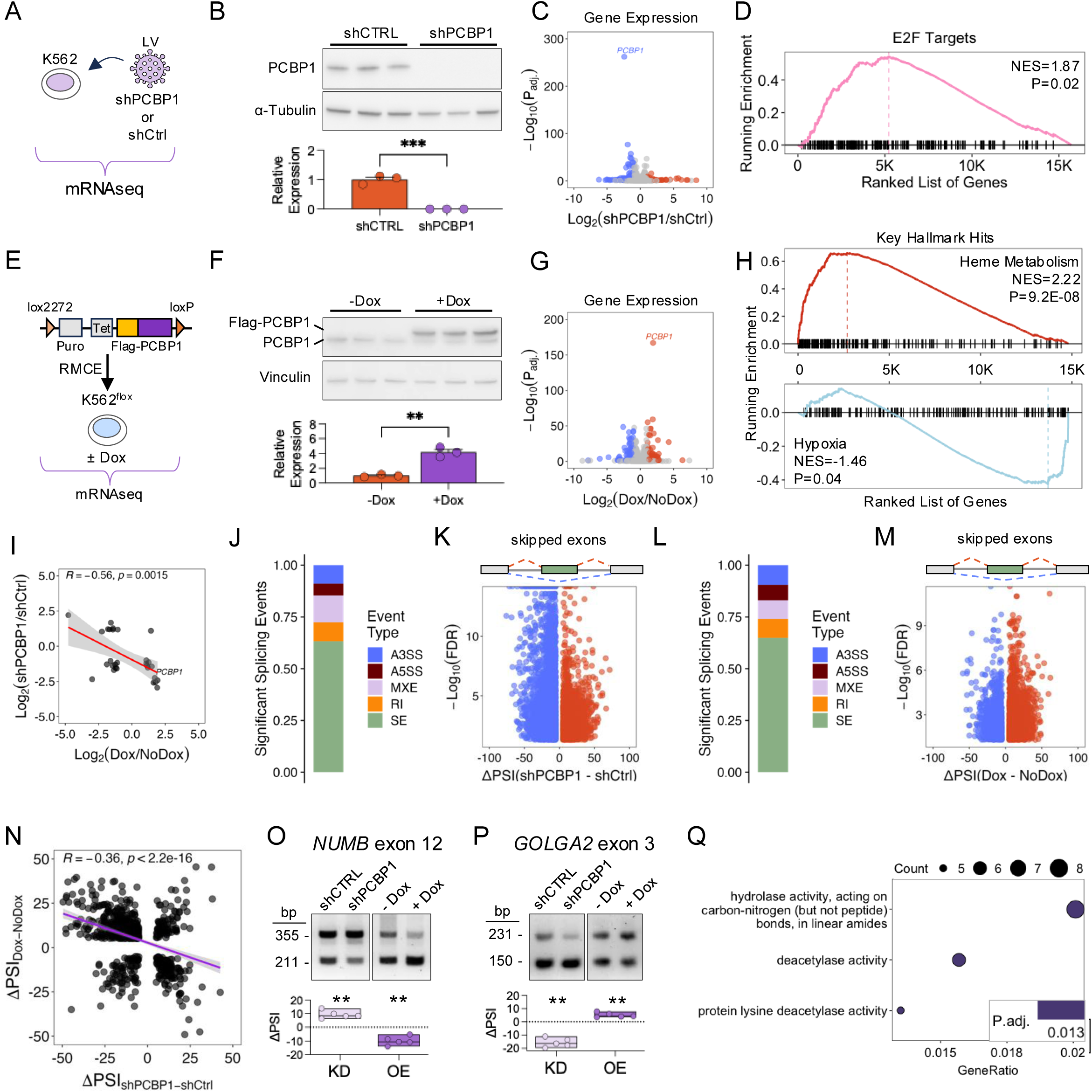
**A.** Schematic of the mRNA-seq experiment of K562 cells transduced with shPCBP1 or shCtrl. **B.** Western blot and quantification of PCBP1 after knockdown. Tubulin was used as a loading control. **B.** Volcano plot of DEGs in shPCBP1 vs. shCtrl. Blue points represent down-regulated genes, and red points represent up-regulated genes. **D.** Enrichment of hallmark ‘E2F targets’ after PCBP1 knockdown by GSEA. **E.** Schematic of the mRNA-seq experiment of K562 cells with doxycycline (Dox)-inducible flag-PCBP1 overexpression. **F.** Western blot and quantification of +Dox and -Dox for Flag-PCBP1 and endogenous PCBP1. Vinculin was used as a loading control. **G.** Volcano plot of DEGs of +Dox versus -Dox. Blue points represent down-regulated DEGs, and red points represent up-regulated DEGs. **H.** GSEA of key hallmark hits after PCBP1 overexpression. **I.** Scatter plot of overlapping significant DEGs between PCBP1 knockdown and PCBP1 overexpression. **J.** Bar plot of the proportion of significant AS events in shPCBP1 vs. shCtrl for each event type. **K.** Volcano plot of SEs with PCBP1 Knockdown. **L.** Barplot of the proportion of significant AS events in PCBP1 overexpression for each event type. **M.** Volcano plot of SEs for PCBP1 overexpression. **N.** Scatter plot of overlapping significant SEs between PCBP1 overexpression and PCBP1 knockdown. **O.** RT-PCR analysis and quantification of *NUMB* exon 12 after PCBP1 knockdown or overexpression. **P.** RT-PCR analysis and quantification of *GOLGA2* exon 3 after PCBP1 knockdown or overexpression. **Q.** GO analysis of genes with opposing of inclusion between PCBP1 knockdown and overexpression. \*\**P*< 0.01, \*\*\**P*<0.001.

As a splicing factor, loss or overexpression of PCBP1 caused many changes in alternative splicing, especially skipped exons (Fig. 2J-M and Supplemental Fig. S2A-H). Gene ontology analysis of alternatively spliced genes after PCBP1 knockdown or overexpression showed shared pathways such as cilium assembly and double strand break repair (Supplemental Fig. S2I-J), suggesting these pathways may be directly regulated by PCBP1. After PCBP1 knockdown, changes in skipped exons accounted for more than 50% of significant alternative events, with more exons skipped than included (4177 skipped and 2218 included, *P* < 2.2e-16, Fig. 2K). Likewise, PCBP1 overexpression also preferentially impacted skipped exons (Fig. 2L and Supplemental Fig. S2E-H) but unlike PCBP1 knockdown, we saw more exon inclusion than exclusion with PCBP1 overexpression (1233 skipped and 2126 included, *P* < 2.2e-16, Fig. 2M). Together, these data indicate that PCBP1 primarily promotes exon inclusion, as shown previously^27^.

Comparing significant skipped exon events across PCBP1 knockdown and overexpression datasets, we found 725 overlapping events, which was found to be significant (*P* < 6e-184). Amongst the overlapping events, about 77% (555) occurred with opposing inclusion/exclusion depending on depletion or over-expression (Fig. 2N), resulting in an overall negative correlation (R=-0.36, *P* < 2.2e-16). We validated skipped exon events in *NUMB* and *GOLGA2* mRNAs. Exon 12 of *NUMB* is more included with PCBP1 knockdown whereas it is more skipped with overexpression (Fig. 2O). In contrast, *GOLGA2* exon 3 is more skipped with PCBP1 knockdown and more included with PCBP1 overexpression (Fig. 2P). NUMB is an endocytic adaptor protein and mutations in a NUMB-associated complex member causes iron accumulation and neurodegeneration in humans^52, 53^. GOLGA2, also known as GM130, is a structural protein involved in the Golgi apparatus and chromosome segregation^54, 55^. More globally, these 725 skipped exon events, likely core PCBP1-regulated splicing targets in K562 cells, are enriched in genes associated with deacetylase activity (Fig. 2Q). These genes included histone deacetylase and sirtuin family members and both groups have been implicated in control of iron homeostasis^56, 57^.

### Overlap between iron-sensitive and PCBP-directed transcriptomes

To assess the iron sensitivity of PCBP-dependent RNA regulation, we integrated changes in gene expression and alternative splicing between iron chelation and PCBP1 knockdown or overexpression. There was a significant overlap in differential gene expression between iron chelation and PCBP1 knockdown, and interestingly, most of these genes were up-or down-regulated in opposing directions (Fig. 3A). We similarly found a significant overlap between differentially expressed genes with iron chelation and PCBP1 overexpression (Fig. 3B). While many genes were up- or down-regulated in opposing directions, there was a greater proportion of genes changing in expression in the same direction. These data suggest that, compared to knockdown, PCBP1 overexpression better mimics gene expression changes with iron chelation raising the possibility that iron negatively impacts PCBP1-mediated control of gene expression.

**Figure 3.**
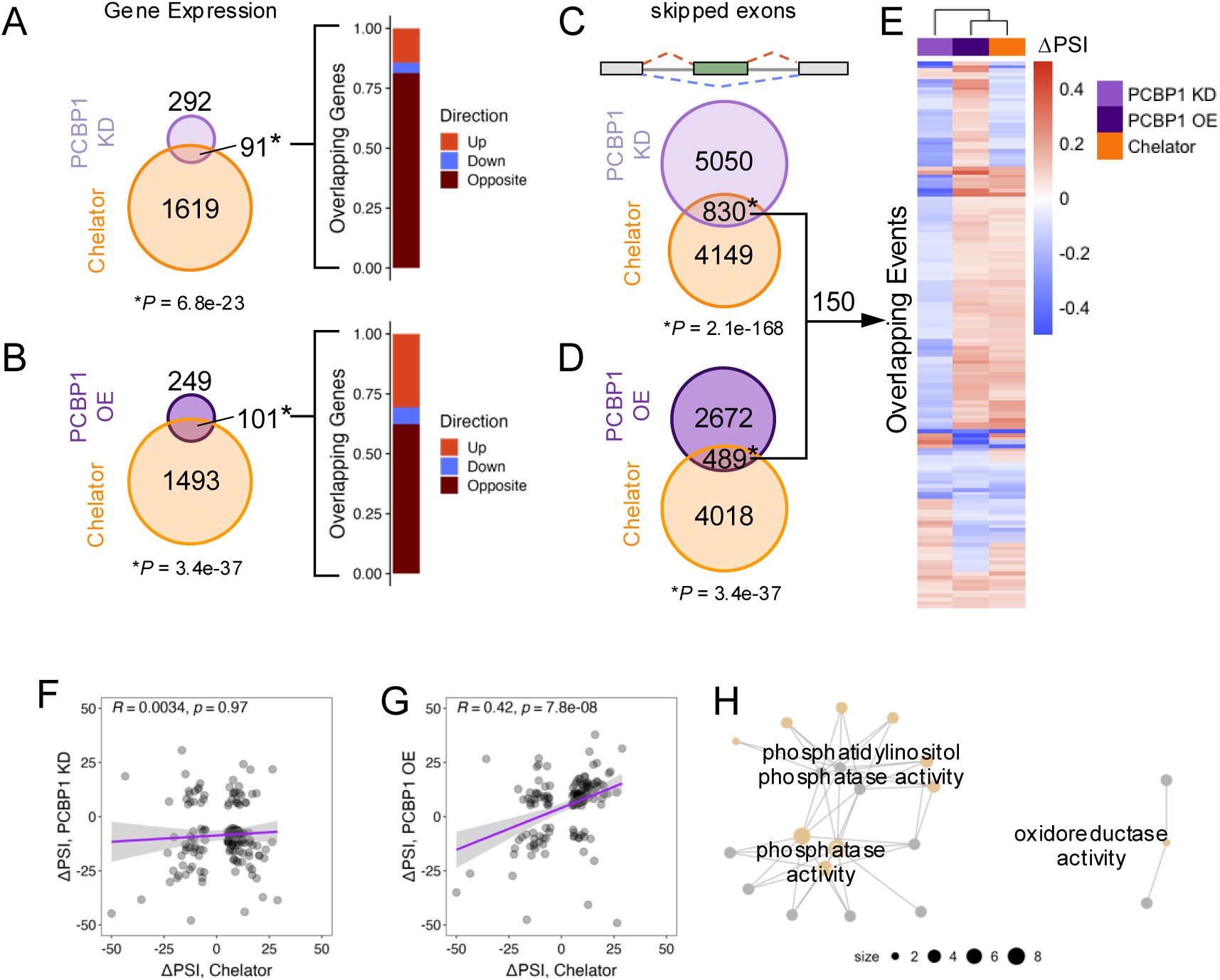
**A.** Venn diagram of significantly DEGs in chelation and PCBP1 knockdown (KD) and bar plot of overlapping genes with direction of regulation (up-regulated, red; down-regulated, blue; or opposing regulation, brown). **B**. Venn diagram comparing significantly DEGs in chelation and PCBP1 overexpression (OE) and bar plot of overlapping genes with direction of regulation (up-regulated, red; down-regulated, blue; or opposing regulation, brown). **C.** Venn diagram comparing significant SEs in Chelator and PCBP1 KD. **D.** Venn diagram comparing significant SEs in Chelator and PCBP1 OE. **E.** Heatmap with clustering of significant SEs between Chelator, PCBP1 KD, and PCBP1 OE. Red is exon inclusion and blue is exon exclusion. **F.** Scatter plot of significant SEs between Chelator and PCBP1 KD from panel E. **G.** Scatter plot of overlapping significant SEs between Chelator and PCBP1 OE from panel E. **H.** Network plot of GO terms from overlapping AS genes between Chelator, PCBP1 KD, and PCBP1 OE.

Next, we examined significantly skipped exon events between iron chelation and PCBP1 knockdown or overexpression. We found a significant overlap in skipped exon events between PCBP1 knockdown and iron chelation (830 exons, Fig. 3C). Similarly, we found a significant overlap in events between PCBP1 overexpression and iron chelation (489 exons, Fig. 3D). In overlapping skipped exon events between iron chelation, PCBP1 knockdown, and overexpression, we found 150 skipped exon events (Supplemental Table 8). We next evaluated the specific inclusion or exclusion (Δ percent spliced in [ΔPSI]) of all 150 events in all three datasets and found that PCBP1 overexpression clustered best with iron chelation (Fig. 3E). Next, we examined the correlation of the ΔPSI of the 150 events between each pair of datasets. There was no significant correlation in skipped exon usage between PCBP1 knockdown and iron chelation (Fig. 3F).

Consistent with splicing between iron chelation and PCBP1 overexpression clustering together, there was a significant positive correlation in skipped exon usage between PCBP1 overexpression and iron chelation (Fig. 3G). Thus, if PCBP1 overexpression promoted exon inclusion, so too did iron chelation. These splicing data mirror our observation of similarities in differential gene expression between PCBP1 overexpression and iron chelation and further support the idea that iron negatively impacts PCBP1 RNA processing. Gene ontology analysis of the 150 overlapping skipped exons revealed a focus on phosphatases, especially phosphatidylinositol phosphatases, and oxidoreductases (Fig. 3H).

To further evaluate the connection between iron chelation and the PCBP-directed transcriptome, we examined PCBP2 knockdown in K562 cells from the ENCODE consortium ^21^. We detected 560 significantly differentially expressed genes (Supplemental Fig. S3A and Supplemental Table 7) involved in catabolic processes and metabolism (Supplemental Fig. S3B). GSEA found negative enrichment for heme metabolism (Supplemental Fig. S3C) and oxidative phosphorylation, among others (Supplemental Table 2). Compared to PCBP1, the loss of PCBP2 caused a notable increase in retained introns (Supplemental Fig. S3D,H), although many other splicing changes were detected (Supplemental Fig. S3D-I). Among skipped exons, we found a significant overlap between iron chelation and shPCBP2 (Supplemental Fig. S3K), further suggesting that PCBP-regulated exons are differentially spliced when iron levels are low. As above, we compared the inclusion or exclusion level for skipped exons under iron chelation versus PCBP2 knockdown. For overlapping events, we found a noteworthy propensity for opposing exon inclusion between iron chelation and PCBP2 knockdown (Supplemental Fig. S3L-M). For example, if an exon was more included with iron chelation, it was likely excluded with PCBP2 knockdown.

Taken together, these data support the intriguing possibility that intracellular iron levels alter the RNA regulatory activity of PCBPs. Specifically, these data suggest that low levels of iron promote PCBP RNA regulation, whereas iron may reduce it, perhaps through reduced RNA binding.

### Low iron levels promote PCBP1 association with RNA in cells

To evaluate how RNA binding by PCBP1 is altered by low iron availability in cells, we turned to a quantitative UV crosslinking and immunoprecipitation method (easyCLIP^58^) (Fig. 4A).

**Figure 4.**
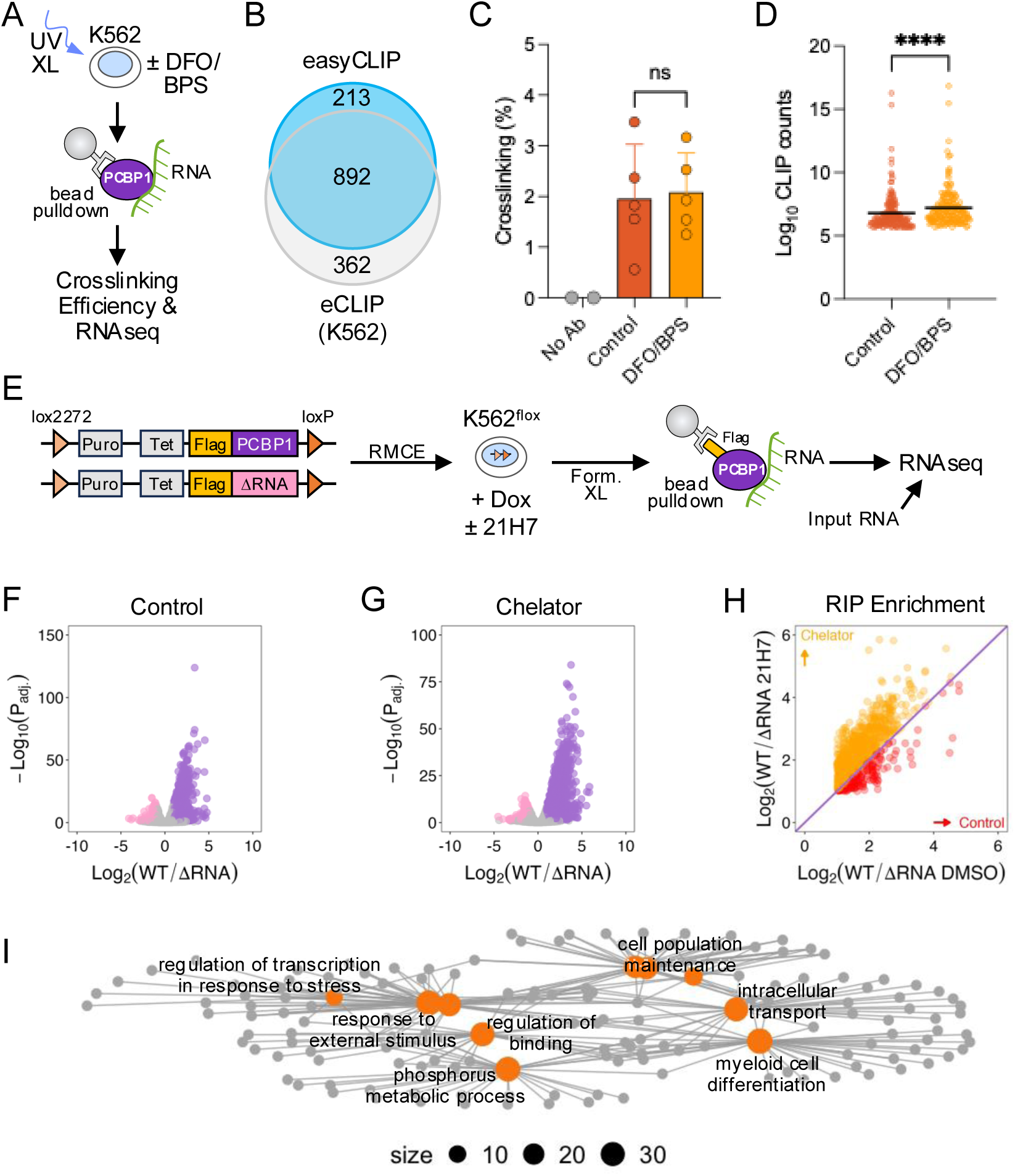
**A.** Schematic of the easyCLIP experiment. **B.** Venn diagram of genes to PCBP1 by eCLIP (ENCODE) and easyCLIP. **C.** Rate of RNA crosslinking to PCBP1 in cells treated with or without iron chelators (DFO/BPS). Data are mean ± SEM. **D.** Dot plot of read counts for genes bound to PCBP1 after control or DFO/BPS treatment. Line is the mean, \*\*\*\**P*<0.0001. **E.** Schematic of RIPseq experiment using the flag-tagged WT or ΔRNA PCBP1. **F.** Volcano plot of genes significantly bound for PCBP1 WT vs. ΔRNA in DMSO treatment. Pink represents genes better bound to ΔRNA PCBP1, and purple represents genes bound better to WT PCBP1. **G.** Volcano plot of genes significantly bound for PCBP1 WT vs. ΔRNA with iron chelation. **H.** Scatter plot of significantly bound genes to WT PCBP1 with iron chelator or DMSO, respectively. **I.** Network plot of genes better bound to PCBP1 after iron chelation.

As with other crosslinking and immunoprecipitation approaches (such as eCLIP), easyCLIP involves UV crosslinking (where only direct protein-RNA interactions are captured), immunoprecipitation of a target RBP and isolation of the bound RNA followed by library preparation. However, easyCLIP also incorporates a method to quantify how much RNA is crosslinked to protein and, therefore, to calculate rate of crosslinking. We performed easyCLIP with K562 cells treated with or without iron chelators (DFO and BPS). We sequenced the bound RNAs and validated our hits, at the gene level, against the ENCODE K562 PCBP1 eCLIP experiment. We found a significant and majority overlap between easyCLIP targets and eCLIP targets (Fig. 4B). Next, we determined the rate of crosslinking to PCBP1 in cells treated with or without iron chelation. Compared to the control, there was no difference with iron chelation in overall RNA crosslinking rate to PCBP1 (Fig. 4C). However, with low iron there was an increase in the association of RNA with PCBP1, on a per gene basis (Fig. 4D). These data suggest PCBP1 has enhanced binding to at least a subset of target RNAs.

Because PCBP1, compared to other RBPs, has an overall low UV crosslinking rate^58^, we turned to formaldehyde crosslinking followed by RNA-immunoprecipitation and sequencing (RIPseq). This approach captures both direct and indirect RNA-protein interactions. For this experiment, we used our doxycycline-inducible flag-tagged wildtype PCBP1 cell line and similarly generated K562 with inducible expression of mutant PCBP1 unable to bind RNA (ΔRNA). These cell lines were treated with either DMSO or iron chelation, followed by formaldehyde treatment, immunoprecipitation for the flag-tagged proteins, and isolation of bound RNA (Fig. 4E). As expected, wildtype PCBP1 bound significantly more genes that ΔRNA PCBP1, in both the control and iron chelator treatments (Fig 4F-G). However, we observed more genes bound to PCBP1 under iron chelation than in the control (2005 versus 1220). Comparing significantly bound genes common between our DMSO and iron chelation data sets, we found 1041 genes, most of which had higher log2 fold changes with iron chelation treatment than control (Fig. 4H). Consistent with pathways observed globally changing with iron chelation (Fig. 1), RNAs bound more to PCBP1 with low iron were involved in myeloid differentiation, DNA regulation, and metabolic processes (Fig. 4I). These data support a model in which low iron levels promote PCBP1 association with RNA, however the mechanism driving this is yet unknown.

### Enhanced PCBP1 binding to RNA with low iron promotes exon inclusion

We next evaluated whether enhanced PCBP1 RNA binding promoted PCBP1-mediated alternative splicing. To do this we compared PCBP1 RIP enrichment (log2 fold change) with or without iron chelation for genes with skipped exons that were either excluded or included more after knockdown or overexpression of PCBP1 (as shown above). First, we looked at skipped exons that were sensitive to loss of PCBP1 and found that both excluded and included exons had increased PCBP1 RIP enrichment with iron chelation compared to control (Fig. 5A). Next, we looked at skipped exons sensitive to PCBP1 overexpression and found that only included exons had enhanced PCBP1 RNA association with iron chelation (Fig. 5B). These data are consistent with PCBP1 as an alternative exon enhancer. Finally, when looking at skipped exons that change with iron chelation (from Fig. 1K), we find increased PCBP1 RNA association with iron chelation for both excluded and included exons (Fig. 5C). These data indicate how low iron levels promote PCBP1 RNA association and enhance alternative splicing control (Fig. 5D). Thus, PCBP iron-sensitive RNA regulation impacts signaling, metabolism, and differentiation through alterative splicing and control of gene expression.

**Figure 5.**
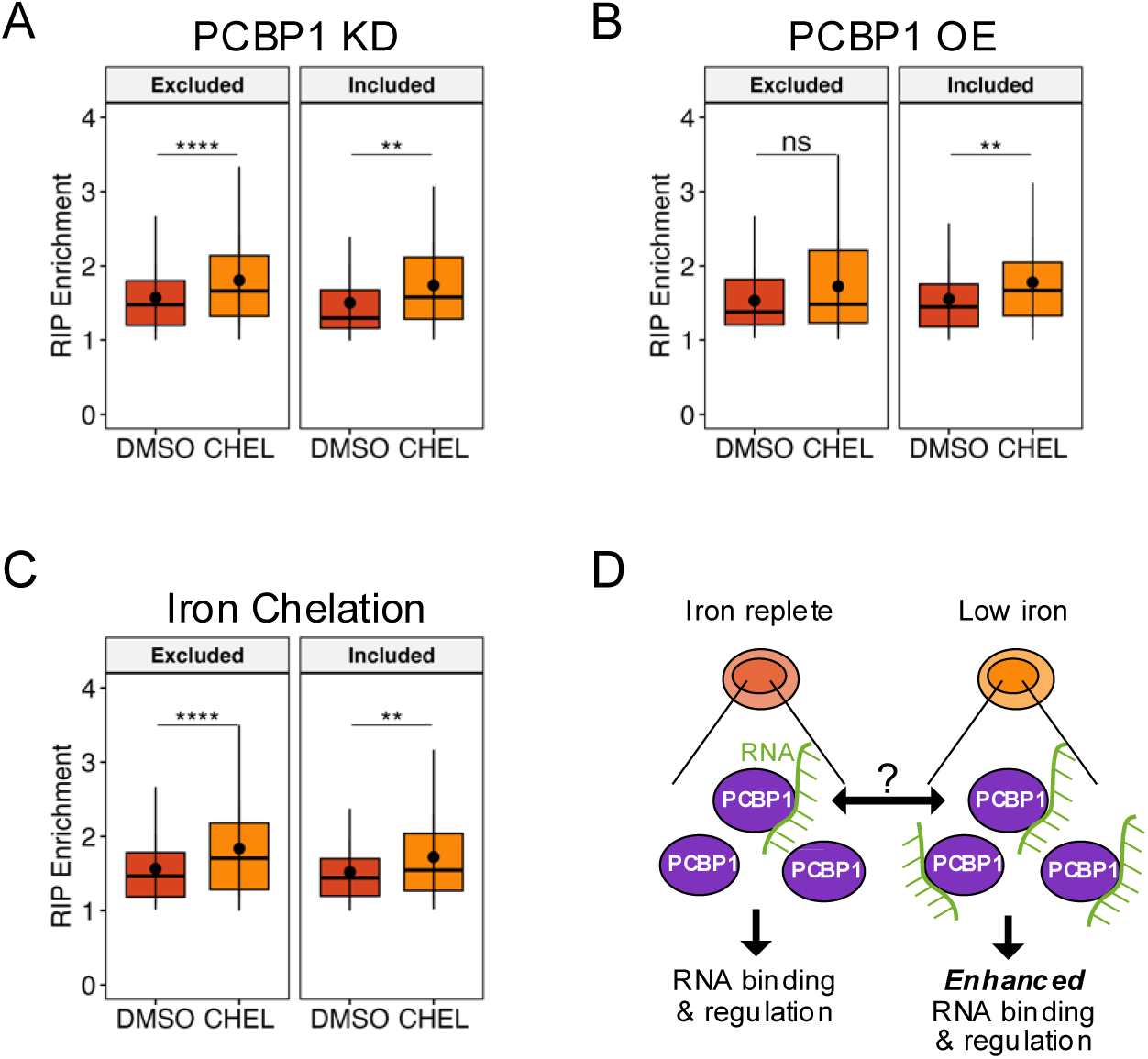
**A.** Box plot of PCBP1 RIP enrichment (with or without iron chelation) for genes with alternatively spliced exons (excluded, left and included, right) after PCBP1 knockdown. **B.** Boxplot of PCBP1 RIP enrichment (with or without iron chelation) for genes with alternatively spliced exons (excluded, left and included, right) after PCBP1 overexpression **C.** Boxplot of PCBP1 RIP enrichment (with or without iron chelation) for genes with alternatively spliced exons (excluded, left and included, right) after iron chelation. **D.** Proposed model of PCBP1 RNA binding and regulation in the presence and absence of iron in cells.

## DISCUSSION

Here we have shown how low iron availability alters the PCBP-directed transcriptome. This finding highlights the variety of posttranscriptional mechanisms available for cells to fine tune their response to stresses like low iron. Via the ability of PCBPs to target thousands of RNAs and regulate their splicing, translation, and/or stability, PCBPs can elicit broad iron-sensitive changes across the transcriptome.

In our study, the PCBP-directed transcriptome hits transcription factor pathways (*e.g.,* E2F), DNA repair and metabolism pathways, signaling pathways, catabolic processes and multiple iron or oxygen-related pathways (*e.g.*, heme metabolism and hypoxia). PCBPs are therefore situated at the nexus of gene expression and metabolism. When we drilled down to only the significant alternative exon changes common across iron chelation and PCBP1 knockdown or overexpression, we found a focus on phosphatidylinositol phosphatases. These metabolic enzymes are key regulators of lipid signaling and have roles in growth factor responses, vesicle transport and cell motility^59^. Ferroptosis, a form of cell death driven by iron-induced lipid peroxidation, is repressed by PCBP1 RNA regulation of specific genes, *BECN1* and *ALOX15*^60^. Perhaps PCBP1 directed alternative splicing of phosphatidylinositol phosphatases contributes to its ability to repress ferroptosis.

Like other RBPs, PCBPs perform their RNA regulatory function as components of ribonucleoprotein complexes. Indeed, PCBP1 interacts with U2AF2 and the U2snRNP at 3’ splice sites of alternative exons^27^. PCBP recruitment of PABP to 3’ untranslated regions mediates mRNA stability^61^. Although there is evidence of PCBP1 iron-sensitive protein-protein interactions^11^, the entire interactome of PCBPs, particularly on RNA, is unclear. More research is needed to clarify how changing iron levels may impact PCBP1 ribonucleoprotein complexes. It is plausible that iron-sensitive changes in PCBP1 protein interactions alters PCBP1-RNA interactions (*e.g.*, PCBP1 free of protein interactors may more readily bind RNA). Iron-sensitive changes in protein-protein interactions could partially explain the differences between easyCLIP and RIP.

PCBPs also control translation of both cellular and viral mRNAs^19^. In particular, PCBP1 functions as a translational repressor^62, 63^. Notably, translation is globally repressed during iron deficiency^64^. Given our finding that reduced iron availability enhances PCBP1 RNA interactions, PCBPs could similarly contribute to global translational repression during iron deficiency.

Our study has several limitations. First, we used an iron chelator to lower available iron levels. Although commonly employed in this fashion, it is possible this approach does not model iron deficiency well and the chelator-iron complex may have off target effects. However, it is of note that iron chelators, like deferoxamine or deferiprone, are used clinically for the treatment of iron overload such as in transfusion-dependent patients with β-thalassemia. These patients primarily present with cardiac iron overload^65^. It is possible, therefore, that iron chelators enter cells not directly experiencing iron overload to produce transcriptome changes as seen here.

Another limitation is our use of a cancer cell line, K562, with known genomic anomalies^66^ that likely influence the transcriptome and metabolism. However, our data from this cell line can be widely integrated with numerous other studies using K562 cells and will be used to generate hypotheses for testing in normal cells or tissues.

Finally, we have not uncovered the specific mechanism for enhanced PCBP RNA interactions with iron chelation. Previous work has demonstrated PCBP1 iron-dependent interactions with BOLA2 that are required for downstream iron-sulfur cluster formation^11^. It is highly probable that low iron levels can drive global changes in PCBP protein-protein interactions. Indeed, one can imagine iron-less PCBPs binding more to RNA simply due to fewer interactions with proteins normally metallated by PCBPs. Another possibility for iron-sensitive RNA binding by PCBP1 is changes in post-translational modifications. For example, PCBP1 is phosphorylated at several sites and thus translocated into or out of the nucleus ^63, 67^. The activity of some phosphatases is iron-dependent (*e.g.*, protein phosphatase 1α^68^) suggesting the possibility that PCBP1 phosphorylation status may be sensitive to cellular iron levels. However, phosphatases that regulate PCBP1 phosphorylation have not yet been identified.

In summary, PCBPs are ubiquitously expressed RBPs that regulate the fate of many RNAs and are essential for life. Their roles as iron chaperones enables them to be highly functionally sensitive to iron levels. Our data suggest PCBPs can integrate iron sensing into changes to the transcriptome. Further work is needed to uncover the likely multifactorial mechanisms that confer this ability.

## METHODS

### Cell Culture and Treatments

K562 cells, including stable cell lines described below, were cultured in RPMI 1640 (Gibco, #11875-093) with 10% fetal bovine serum (FBS, Corning, #35-015-CV) and 1x Anti-Anti (Gibco, #15240062) and incubated at 37° C with 5% CO_2_. Cultures were periodically tested for mycoplasma contamination with MycoStrip tests (Invivogen, #rep-mysnc-50). For iron chelation experiments, cells were treated with 10 µM 21H7 or 100% DMSO (volume-matched) in RPMI 1640 complete media overnight. K562 cells used in easyCLIP experiments (described below) were treated overnight with 100 µM deferoxamine mesylate (DFO, Sigma, #252750) and 100 µM bathophenanthroline disulfonic acid (BPS, Sigma, #146617) in RPMI 160 complete media.

### Lentiviral Transduction for Gene Knockdown

Lentiviral particles containing pLKO.1 plasmids with shRNA genes for Luciferase (shCtrl) or PCBP1 (shPCBP1) were generated as described from HEK293T cells^69, 70^. K562 cells were treated with media containing lentiviral particles and 8µg/mL polybrene, overnight at 37° C with 5% CO_2_. The following morning, cells were spun down (500 *x g*, 3 min) and resuspended in selection media (3 µg/mL puromycin, 10% FBS, RPMI 1640) for four days before collection.

### Generation of K562^flox^ cells

K562 cells were electroporated using the Neon^TM^ Transfection System (Thermo Scientific, #MPK1025) with AAVS1 guide RNA (Synthego), Cas9-NLS (Synthego), and a donor plasmid (generous gift from Matt Taliaferro, Univ. of Colorado Anschultz) containing a blasticidin-resistance cassette flanked by loxP and lox2272 sites under an EF-1α promotor with homology regions to the AAV1 locus that is compatible with the pRD-RIPE system for recombinase-mediated cassette exchange (RMCE) ^51^. Cells were selected with 10 µg/mL blasticidin (MilliporeSigma, # SBR00022) and plated for single cell clonal expansion. The final clone was genotyped using sanger sequencing.

### Generation of stable cells lines with inducible gene expression in K562^flox^

K562^flox^ cells were electroporated using the Neon^TM^ Transfection System with pRD-Cre and pRD-RIPE plasmids (generous gift from Eugene Makeyev, King’s College London) modified to contain Flag-PCBP1 or Flag-PCBP1-ΔRNA (R40A, R124A, and R306A as in ^22^) and incubated overnight in RPMI 1640 with 10% FBS in an incubator at 37 °C and 5% CO_2_. The next day, puromycin (1 µg/mL, final, Thermo Scientific, #A1113803) was added to culture media for selection. Polyclonal stable cell lines were maintained and expanded under puromycin selection for downstream experiments.

### Western Blotting

Normalized cell lysates prepared in 1x RIPA buffer (Sigma-Aldrich, #20-188) with 1x Halt Protease Inhibitor Cocktail (Thermo Fisher, #78420) were submitted to SDS-PAGE and transferred to nitrocellulose membranes. Membranes were blocked with 5% milk or 5% bovine serum albumin (BSA, Fisher, #BP9706100) for 30 minutes at room temperature. Primary antibodies were incubated overnight at 4° C then washed repeatedly with 1x TBST buffer (Tris-buffered saline, 0.05% Tween-20, Thermo Fisher, #J77500.K2). Blots were incubated with peroxidase-conjugated secondary antibodies for 30 minutes to 1 hour at room temperature followed by repeated TBST washes. Blots were then incubated with up to 4-fold diluted western ECL substrate (Bio-Rad, # 1705061) for up to 5 minutes. Blot chemiluminescence was imaged with a ChemiDoc imager (Bio-Rad). Band intensities were measured with ImageJ software.

Antibodies used are as follows: Rabbit anti-PCBP1 polyclonal (1:2000, MBL International Corp, #RN024P), Rabbit anti-FTH1 polyclonal (1:1000, Cell Signaling, #3998S), Rabbit anti-IRP2 (D6E6W) monoclonal (1:1000, Cell Signaling, #37135S), Rabbit anti-α-Tubulin polyclonal (1:5000, Cell Signaling, #2144S), Rabbit anti-Vinculin polyclonal (1:5000, Proteintech, #26520-1-AP), Goat anti-Rabbit-HRP-conjugated polyclonal (1:10,000, Jackson ImmunoResearch, #111035045), Donkey anti-Rabbit-HRP-conjugated polyclonal (1:10,000, Jackson ImmunoResearch, #711035152).

### mRNAseq

RNA isolated from cells after treatments was sent to Novogene (Sacramento, CA) for polyA selection and library preparation followed by Illumina sequencing (paired end, 150 cycles). Eighteen to 30 GB of data per sample were collected.

### mRNAseq Analysis

Quality of sequencing fastq files was checked using FastQC^71^ and MultiQC^72^. Salmon (v1.10.2)^73^ was used for transcript abundance quantification which was then analyzed for differential gene expression using DESeq2 (v3.19)^74^. Significant genes were quantified by passing a read count threshold greater than 10, having an absolute log2 fold change greater than 1 and an adjusted P value less than 0.05. To capture alternative splicing, reads were first mapped to human genome build hg38 from Ensembl^75^ using STAR (v2.7.11b)^76^. The resulting bam files were then analyzed with rMATS^77^. Significant alternative splicing events had an FDR less than 0.05, an absolute ΔPSI > 5 percent, and having either an average inclusion read count or exclusion read count >= 10. Gene Ontology (GO) and Gene Set Enrichment Analysis (GSEA) were performed using functions from the R/Bioconductor package, clusterProfiler ^78^, including enrichGO and GSEA, respectively. GSEA of Hallmark^35^ gene sets was performed on pre-ranked lists of genes with a ranking metric defined as log_2_(FC)*(1-pvalue) and using the msigdbr package in R . Chi-square tests were used to determine significant differences between observed and expected results. Hypergeometric tests were used to calculate the significance of overlapping datasets, where only co-detected genes or events were used as background. Spearman’s rank correlation test was used in correlation analysis.

### Hypoxia RNAseq

mRNAseq data of K562 cells treated with normoxia (21% oxygen) or hypoxia (1% oxygen) for 3 days by Jain et al.^39^ (GEO Accession: GSE144527) was analyzed using GEO2R (NCBI) which computes differential gene expression using DESeq2. Significant genes had adjusted P values < 0.05 and an absolute log2 fold change > 1.

### Splicing Event Validation

Significant alternatively spliced exon events having a FDR less than 0.05, an absolute ΔPSI > 5 percent, and having an average inclusion read count or exclusion read count >= 10, were determined to be a strong splicing event and were further evaluated for: 1) having clear annotation in UCSC Genome Browser^79^, 2) construction of a pair of PCR primers in Primer3web^80^, and 3) using the UCSC In-Silico PCR tool, PCR primers produced between 2 and 3 output files in fasta format that contain all sequences in the database that lie between and include the primer pair^79^.

K562 cells were treated with iron chelation, PCBP1 knockdown or PCBP1 overexpression with 5 biological replicates, as described above. RNA was isolated from treated K562 cells using phenol:chloroform extraction and cDNA was produced using the High-Capacity cDNA Reverse Transcription Kit (Invitrogen, #4368814). Primer oligos were designed targeting flanking exons of the alternative exon and synthesized by Integrated DNA Technologies. PCRs were performed using Phusion polymerase, 1x GC Buffer, 200 µM dNTPs, 0.5 µM forward and reverse primers, 3% DMSO, and 40 ng cDNA with 27 or less cycles. Some PCR products were run on 6% TBE gels (Invitrogen, #EC6865BOX) which were then stained with SYBR-Gold DNA stain (Invitrogen, #S11494) for 5 minutes with shaking before imaging on a ChemiDoc imager. Other PCR products were run on 2 or 3% agarose gels containing Sybr Safe DNA stain (Invitrogen, #S33102) and imaged on a ChemiDoc imager.

### UV crosslinking and immunoprecipitation (easyCLIP)

K562 cells were treated overnight with DFO and BPS as described above (Cell Culture and Treatments). The easyCLIP protocol was followed as described^58^ with the following adjustments. At harvest, K562 cells (∼45 million per condition) were spun at 200 *xg* for 5 minutes, media removed by aspiration, then resuspended in 3 mL 4°C PBS. The PBS/cell mixture was delivered to a 10 cm plate on a bed of ice and UV (254 nm) cross-linked at 400 mJ/cm^2^. After lysis, polyclonal rabbit anti-PCBP1 antibodies (MBL International Corp, #RN024P) were used to immunoprecipitate PCBP1. Bound RNA libraries were prepared and sequenced (paired end, 150 cycles) on a MiSeq machine and analyzed according to previous methods^58^, mapping to the hg38 genome and counting reads per feature with HTSeq.

### Formaldehyde crosslinking and RNA immunoprecipitation (RIPseq)

RIP experiments were performed as described^81^. Expression of Flag-PCBP1 or Flag-PCBP1-ΔRNA in K562^flox^ stable cell lines was achieved with 200 ng/mL doxycycline (dox, MilliporeSigma, #D9891) overnight. Dox-treated cells were also treated with either DMSO (200,000 cells/mL) or 10 µM 21H7 (400,000 cells/mL) overnight. The next day, cells were counted, washed twice in PBS, and rotated for 30 minutes at 4°C in 10 mL of 0.3% formaldehyde (Fisher, #28906) in PBS. Crosslinking was quenched with 200 mM glycine (final, Fisher, #BP381) for 5 minutes, with rotation, at room temperature then washed twice with cold PBS before resuspension at 10 million cells/mL. Cells were pelleted and snap frozen with a dry ice-ethanol bath and stored at -80°C.

Protein A/G PLUS agarose beads (25 µL per cell pellet, Santa Cruz Biotechnology, #sc-2003) were washed three times in blocking buffer (0.5% BSA in 1xPBS), incubated overnight, with rotation, at 4°C temperature with 10 µL M2 anti-Flag antibody (MilliporeSigma, #F1804), and washed three times with fRIP buffer (25 mM Tris-HCl pH 7.5, 0.5% NP-40, 5 mM EDTA pH 8.0 150 mM KCl, 0.5 mM DTT, 1x Halt protease inhibitor cocktail, 1:200 SUPERaseIN) before use. RIPA buffer (50 mM Tris pH 8.0, 1% Triton X-100, 0.5% sodium deoxycholate, 0.1% SDS, 5 mM EDTA pH 8.0, 150 mM KCl, 0.5 mM DTT, 1x Halt protease inhibitor cocktail, 1:200 SUPERaseIN) was added to each cell pellet. Each sample was sonicated twice (30% amplitude, 2 x 30 seconds on). All samples were centrifuged at 4°C for 15 minutes at 15,000 *x g* and supernatant was transferred to a new tube. 50 µL from each sample was pulled as inputs and stored at -80°C. Samples were normalized by concentration of Flag-tagged proteins detected by Western blot. Cell lysates and beads were incubated overnight at 4°C with rotation. Beads were rinsed once with 1 mL cold fRIP buffer, then washed three times in PolII chip buffer (50 mM Tris H-Cl pH 7.5, 140 mM NaCl, 1 mM EDTA pH 8.0, 1 mM EGTA pH 8.0, 1% Triton X-100, 0.1% sodium deoxycholate, 0.1% SDS), twice in high salt PolII chip buffer (50 mM Tris-HCl pH 7.5, 500 mM NaCl, 1 mM EDTA pH 8.0, 1 mM EGTA pH 8.0, 1% Triton X-100, 0.1% sodium deoxycholate, 0.1% SDS), and once in LiCl buffer (20 mM Tris pH 8.0, 0.5% NP-40, 1 mM EDTA pH 8.0, 250 mM LiCl, 0.5% sodium deoxycholate). Beads were rotated 5 minutes at 4°C before centrifugation at 1200 *x g* for 2 min at 4°C for washes. Sample tubes were changed after the first and final wash. After washes, beads were resuspended in 56 µL water and 59 µL of reverse crosslinking buffer (5 mM DTT, 1:5 Proteinase K, 1:100 SUPERaseIN). Samples were incubated at 42°C for 1 hour, 55°C for 1 hour, and 65°C for 30 min. RNA was subsequently extracted with trizol/chloroform and cleaned with the Zymo RNA Clean and Concentrator-5 columns (Zymo Research, #R1013) with DNA digestion.

Library preparation of isolated RNA from Flag-PCBP1 immunoprecipitations or from whole cell lysates (inputs) was performed with the KAPA RNA Hyperprep Kit with RiboErase (Roche, #KK8560) using NEBNext Multiplex Oligos (adaptors and single index primers, New England Biolabs, #E7335, #E7500, #E7730) compatible with Illumina sequencing. Pooled libraries were sequenced (single end, 100 cycles) on an Illumina Nextseq1000. Quality of sequencing fastq files was checked using FastQC^71^ and MultiQC^72^. Salmon (v1.10.2)^73^ was used for transcript abundance quantification which was then analyzed for differentially bound RNAs using DESeq2 (v3.19)^74^ with a Likelihood Ratio test where bound RNAs were controlled for baseline expression by comparison to input samples, as described^82^.

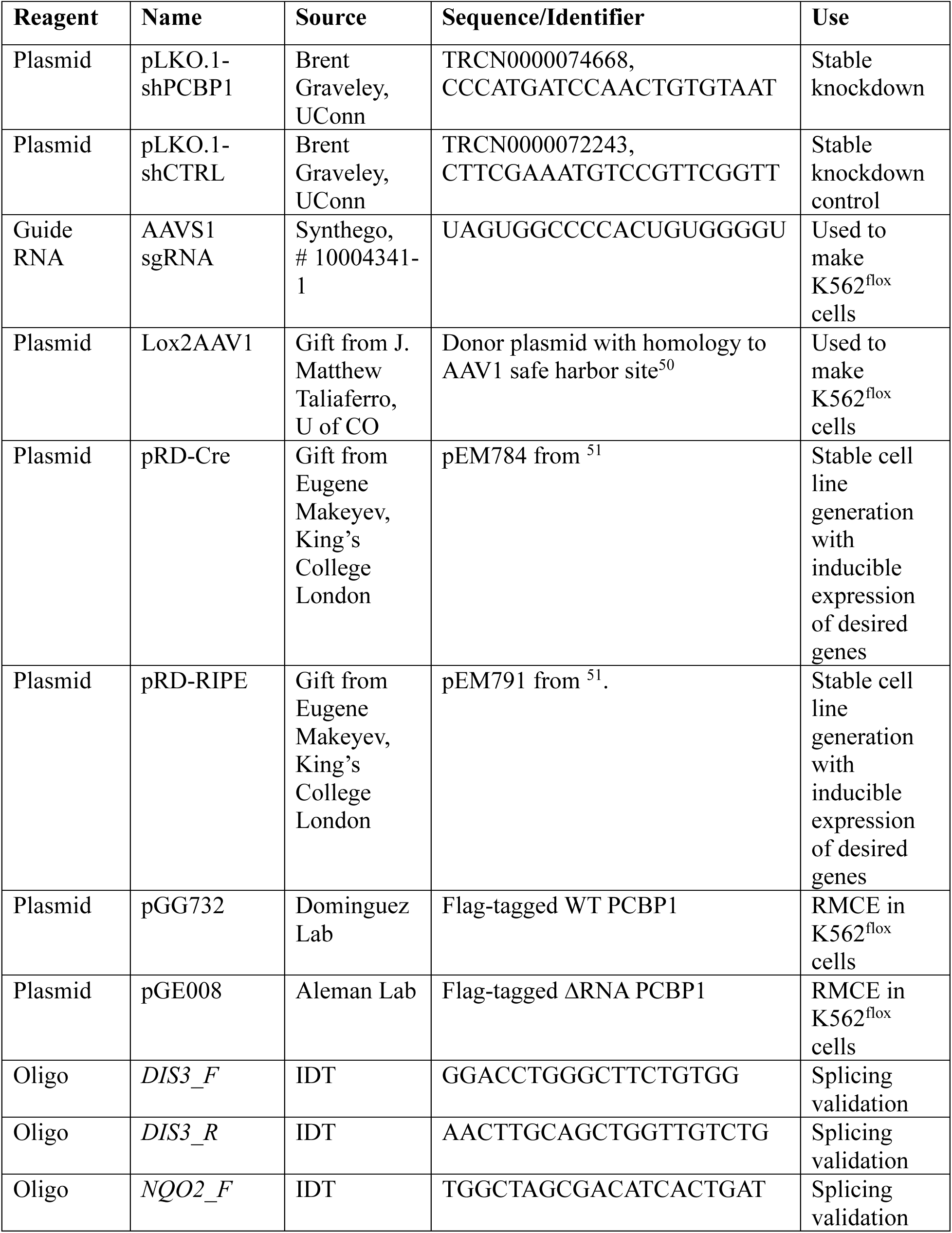
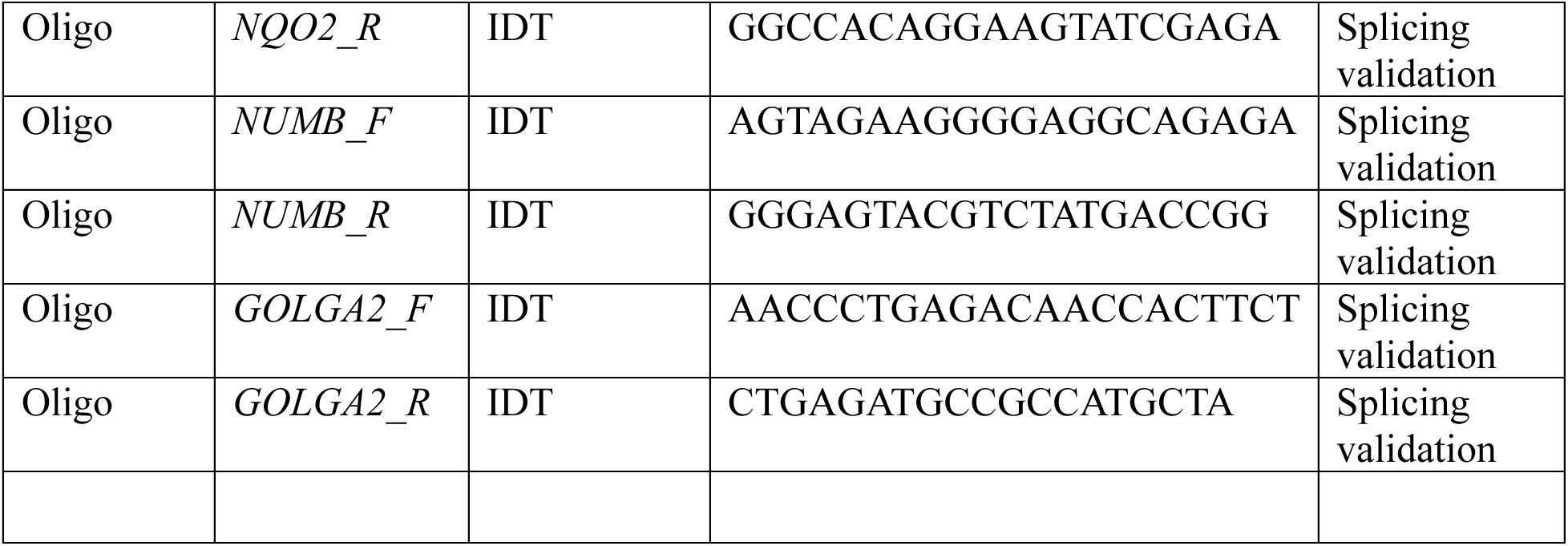
Table: Key Reagents

## Acknowledgements

The authors thank Drs. J. Matthew Taliaferro and Eugene Makeyev for providing plasmids. This study was funded by NIH grants TL1DK139567 (K.F.), R35GM142864 (D.D.) and R01DK124773 (M.M.A.).

---

**Figure S1.**
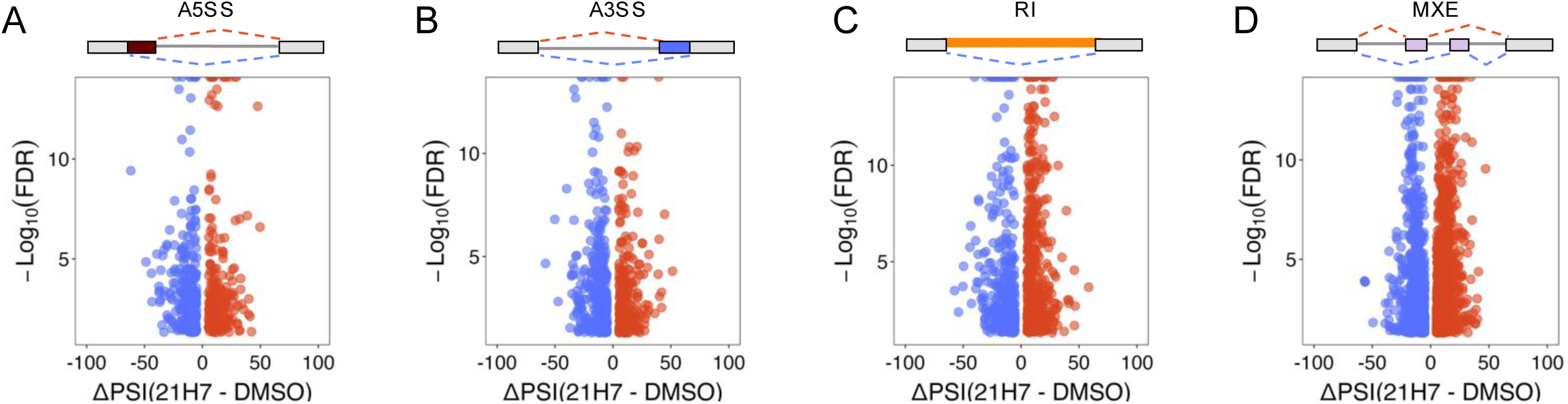
**A-D.** Volcano plot of alternative 5’ splice sites (A5SS), alternative 3’ splice sites (A3SS), retained introns (RI) and mutually exclusive exons (MXE), respectively, for 21H7 vs. DMSO. Blue points are excluded exon events and red points are included exon events.

**Figure S2.**
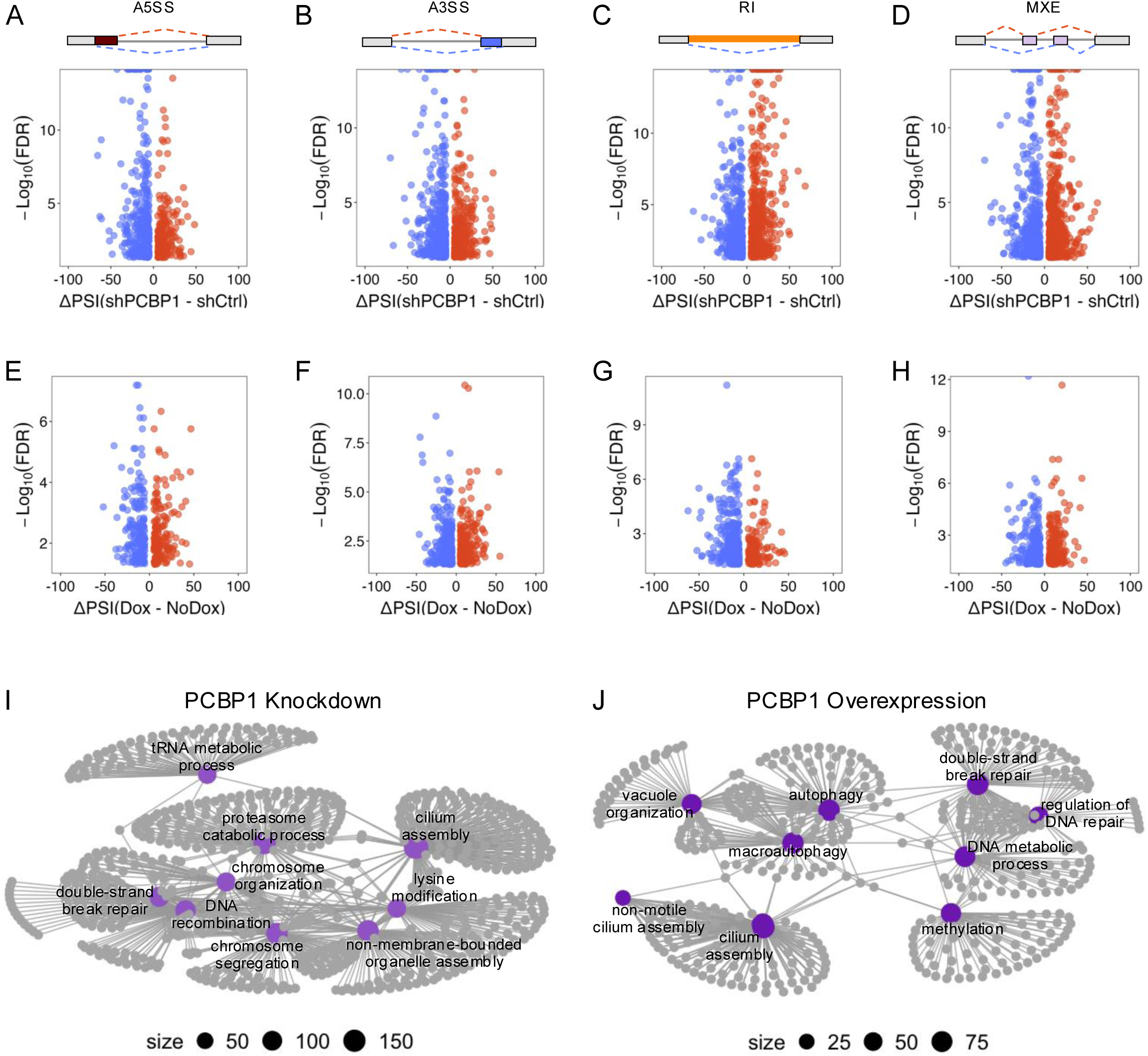
**A-D.** Volcano plots of A5SS, A3SS, RI and MXE, respectively, for shPCBP1 vs. shCtrl. Blue points are excluded exon events, and red points are included exon events. **E-H.** Volcano plot of alternative A5SS, A3SS, RI and MXE, respectively, for PCBP1 overexpression (Dox) vs. control (-Dox). **I.** Network plot of GO terms from significant AS genes from PCBP1 knockdown. **J.** Network plot of GO terms from significant AS genes of PCBP1 overexpression.

**Figure S3.**
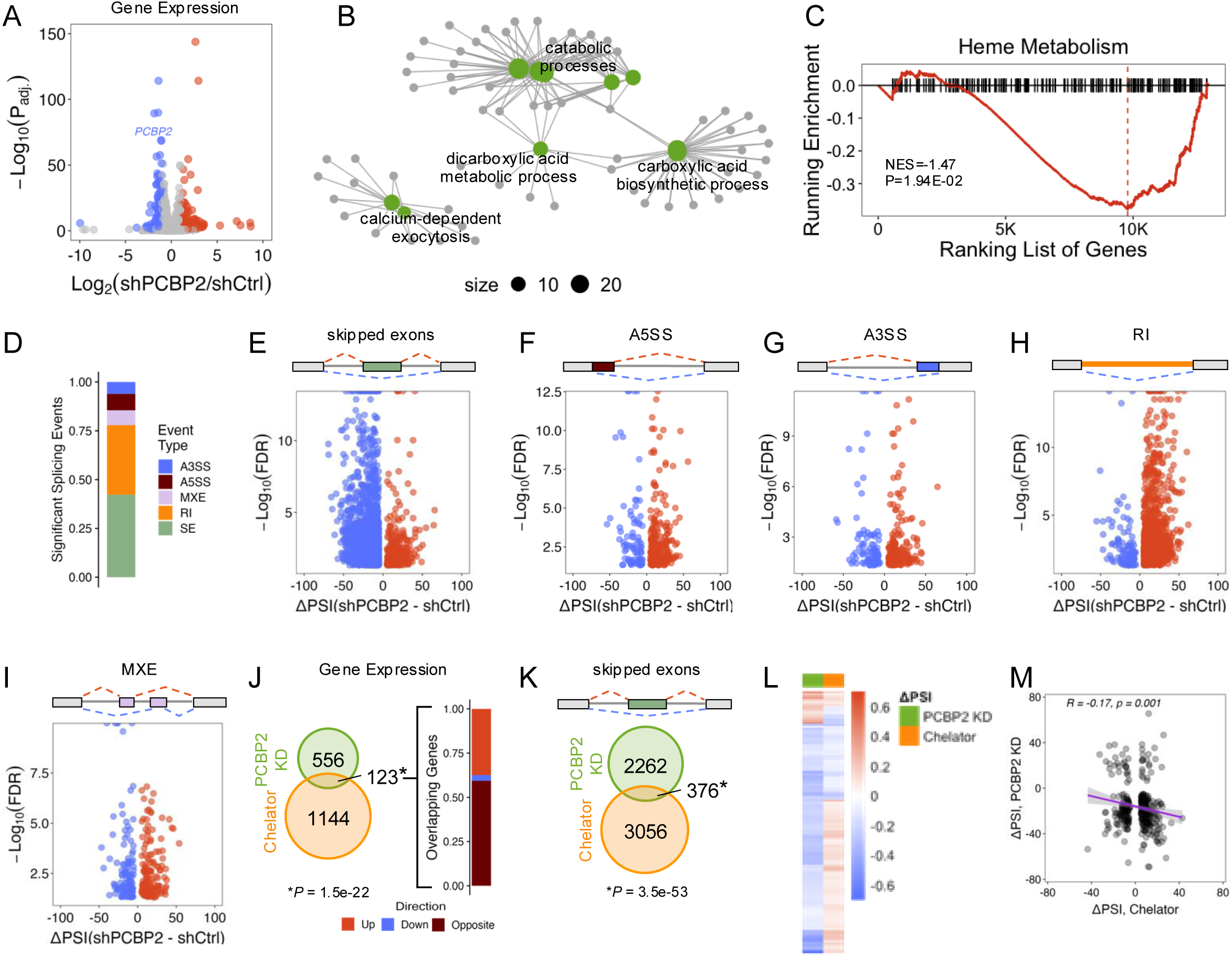
**A.** Volcano plot of DEGs of PCBP2 KD vs. shCtrl. **B.** Network plot of GO terms from significantly DEGs after PCBP2 KD. **C.** Enrichment of hallmark heme metabolism genes after PCBP2 KD. **D.** Bar plot of the proportion of significant AS events in PCBP2 KD vs. shCtrl for each event type. **E-I.** Volcano plots of A5SS, A3SS, RI and MXE, respectively, for PCBP2 KD. **J.** Venn diagram comparing significantly DEGs in Chelator and PCBP2 KD and bar plot of overlapping genes with direction of regulation (up-regulated, red; down-regulated, blue; or opposing regulation, brown). **K.** Venn diagram comparing significant SEs in Chelator and PCBP2 KD. **L.** Heatmap of significant SEs between Chelator, and PCBP2 KD. Red is exon inclusion and blue is exon exclusion. **M.** Scatter plot of significant SEs overlapping between Chelator and PCBP2 KD.

